# Nutrishield^®^ β-Carotene Attenuates LPS-Induced Systemic Inflammation and Oxidative Stress in Murine Model

**DOI:** 10.64898/2026.03.13.711625

**Authors:** Joel Fu, Shaylynn Yu, Kevin Ngo, Xuan Hao Teo, Muhammad Ovais

## Abstract

β-Carotene is a strong antioxidant and immunomodulator, but it breaks down in the presence of oxygen and isn’t readily bioavailable. Nutrishield® β-Carotene is a special starch-based microencapsulation system designed to protect β-carotene from oxidation, make it easier to spread in the gastrointestinal environment, and improve its effectiveness. This study reported Nutrishield® as a formulation platform by using β-carotene as a model compound. We compared how well it worked *in vitro* and *in vivo* with β-carotene that wasn’t encapsulated. We randomly put thirty-two male BALB/c mice into four groups (n=8): Control, LPS, LPS + β-Carotene (10 mg/kg/day), and LPS + Nutrishield® (the same dose of β-Carotene). After an intraperitoneal injection of LPS (1 mg/kg, twice weekly), the treatments were administered orally for 28 days. LPS administration significantly elevated TNF-α (142 ± 8 pg/mL), IL-6 (118 ± 7 pg/mL), and serum 8-OHdG (5.4 ± 0.3 ng/mL) in comparison to controls (TNF-α = 62 ± 5 pg/mL; IL-6 = 51 ± 4 pg/mL; 8-OHdG = 2.1 ± 0.2 ng/mL). Nutrishield® β-Carotene supplementation significantly normalized cytokine profiles (TNF-α = 69 ± 4 pg/mL; IL-6 = 56 ± 3 pg/mL; IL-10 increased from 38 ± 2 to 76 ± 3 pg/mL; p < 0.001 vs. LPS). Histological analysis demonstrated reduced hepatocellular vacuolation and renal tubular degeneration in the Nutrishield® β-Carotene group relative to the β-carotene and LPS groups. These findings demonstrate that Nutrishield® microencapsulation significantly enhances the functional antioxidant and anti-inflammatory efficacy of β-carotene in a murine model of systemic inflammation.

**Graphical Abstract:** 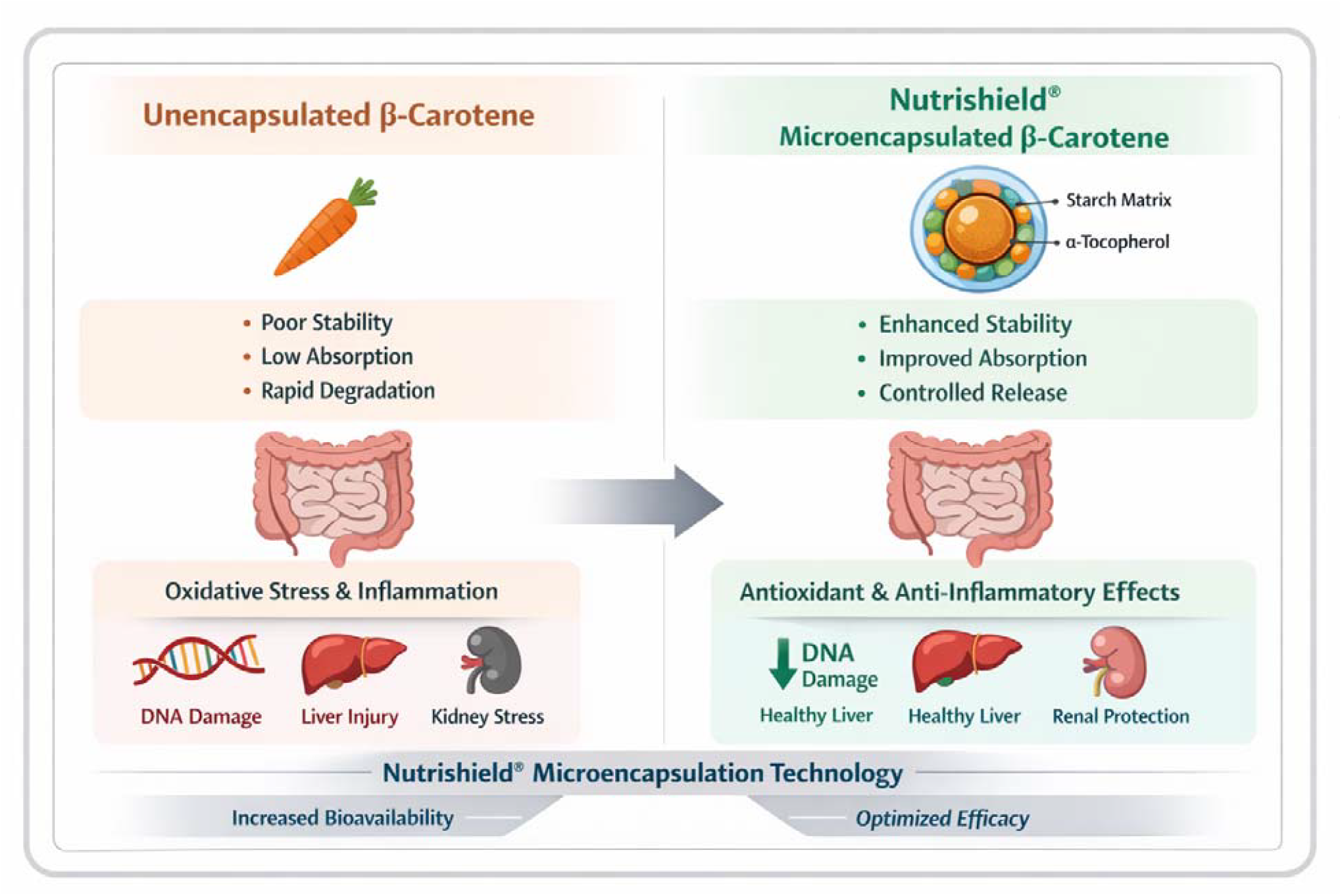

## 1. Introduction

β-Carotene is a carotenoid that dissolves in fat and is well known as a precursor to vitamin A and for its strong antioxidant properties. Carrots, sweet potatoes, and spinach are colorful fruits and vegetables that contain β-carotene. β-carotene supports vision, maintains epithelial cell health, modulates the immune system, and maintains cellular redox balance (Gómez-Mascaraque, Perez-Masiá, González-Barrio, Periago, & López-Rubio, 2017; Jeyakodi, Krishnakumar, & Chellappan, 2018). Because β-carotene is an antioxidant, it can effectively neutralize reactive oxygen species (ROS) and prevent lipid peroxidation. (Zhou et al., 2018).

The medical use of pure β-carotene is limited because it isn’t very bioavailable and breaks down when exposed to oxygen (Bas, 2024). β-carotene can break down during digestion and storage, making it harder for the body to absorb and lowering its levels. Microencapsulation matrices composed of starch not only inhibit the degradation of β-carotene in oxygen-rich environments but also promote its regulated release and enhance intestinal absorption (Donhowe & Kong, 2014). Nutrishield^®^ β-Carotene formulation is developed under the brand name Nano Singapore (https://nanosingaporeshop.com/pages/about-us) by Singapore Ecommerce Centre Pte Ltd, which uses a starch matrix strengthened with all-rac-α-tocopherol (vitamin E) to improve stability against oxidation and mimic the absorption of natural lipids. Nutrishield® is designed to address the physicochemical challenges posed by lipophilic bioactives such as β-carotene. The LPS-induced inflammation model in mice is a well-established paradigm for studying systemic inflammation, mimics bacterial endotoxemia, and triggers the production of pro-inflammatory cytokines such as TNF-α, IL-6, and IL-1β (Skrzypczak-Wiercioch & Sałat, 2022) (Jomova et al., 2023; Leyane, Jere, & Houreld, 2022).

Microencapsulation improves the perceived bioavailability of carotenoids by helping them spread out in the intestinal lumen and mimicking lipid-mediated absorption pathways. Previous research has shown that encapsulation techniques, such as starch-based matrices, lipid carriers, and polysaccharide systems, can enhance carotenoid stability and functional absorption by about 2 to 5 times, without changing the chemical structure. In this context, Nutrishield® serves as a delivery system that maintains β-carotene bioactivity and provides long-lasting antioxidant effects, rather than relying on high doses.

Previous studies have demonstrated that β-carotene diminishes the synthesis of inflammatory cytokines, enhances the function of antioxidant enzymes, and protects against tissue injury in LPS models (Bai et al., 2005; Kawata, Murakami, Suzuki, & Fujisawa, 2018). However, there is a lack of comparative analyses between pure β-carotene and novel formulations such as Nutrishield® β-Carotene. Using an LPS-induced BALB/c mouse model, this study looked at how Nutrishield® β-Carotene compared to pure β-carotene in terms of its anti-inflammatory, antioxidant, and organ-protective effects. A lot of research has been done on β-carotene’s ability to fight free radicals and modulate the immune system, but it’s hard to get consistent biological results because it breaks down when exposed to oxygen and isn’t very bioavailable. Nutrishield® β-Carotene is a formulation-driven solution that uses β-carotene as a model compound to demonstrate the utility of a starch-based microencapsulation platform. This study doesn’t just examine how β-carotene works in the body; it also examines how microencapsulation in the Nutrishield® system makes it more stable, available, and effective during inflammation.

## 2. Materials and Methods

### 2.1. Formulation of Nutrishield^®^ β-Carotene

The microencapsulated beadlets were manufactured using advanced spray-and-starch-capture drying technology (proprietary information provided under NDA). Nutrishield^®^ β-Carotene 20% TAB-S consists of red or reddish-brown free-flowing beadlets, with white spots of food starch. The individual particles containing β-Carotene are finely dispersed in the matrix of modified food starch, coated with corn starch, while all-rac-α-tocopherol is added as an antioxidant. The final composition of Nutrishield^®^ β-Carotene 20% TAB-S is as follows: β-Carotene 20 %, Modified Food Starch 56 %, Corn Starch 21.0%, all-rac-α-Tocopherol 3.0%.

### 2.2. Experimental Animals and Ethics

We obtained 32 male BALB/c mice (6–8 weeks old, 25 ± 2 g) from a licensed animal facility at the National Institute of Health, Islamabad, Pakistan, and acclimated them to their new environment for 1 week (22 ± 2°C, 12 hours of light and 12 hours of dark, and 50–60% humidity). The Institutional Animal Ethics Committee approved all procedures with reference No. 9775, which followed the OECD and ARRIVE 2.0 guidelines.

### 2.3. Experimental Design

Mice were randomized into four groups (n = 8 per group), and all treatments were administered for 28 consecutive days.

| GROUP | TREATMENT DESCRIPTION |
| --- | --- |
| CONTROL | Saline i.p. + Corn oil orally |
| LPS | LPS (1 mg/kg i.p., twice weekly) + Corn oil orally |
| LPS + B-CAROTENE | LPS + β-Carotene (10 mg/kg/day, oral) |
| LPS + NUTRISHIELD® | LPS + Nutrishield® (equivalent to 10 mg/kg β-carotene/day, oral) |

### 2.4. Induction of Inflammation

To cause chronic systemic inflammation, LPS (E. coli 055:B5) from Beijing Solarbio Science & Technology Co., Ltd. was dissolved in sterile saline and injected into the peritoneum (1 mg/kg) on days 1, 4, 8, 12, 16, 20, 24, and 28.

### 2.5. Sample Collection

On day 28, blood samples were taken by puncturing the back of the eye while the mice were lightly sedated with isoflurane. We separated the serum and plasma and put them in a freezer at □80°C. We cut, weighed, and processed liver and kidney tissues for histological or biochemical testing.

### 2.6. Biochemical and Oxidative Stress Assessments

#### Serum Cytokine Quantification (IL-6, TNF-α, IL-10)

We used sandwich ELISA kits from BioLegend (USA) to measure IL-6, TNF-α, and IL-10 levels in serum. We followed the instructions that came with the kits. In short, serum samples (diluted 1:2–1:5 depending on the cytokine concentration) and corresponding standards were added to 96-well plates precoated with capture antibodies. They were then left at room temperature for 2 hours. After washing with PBS-T (0.05% Tween-20), biotinylated detection antibodies were added and incubated for 1 hour. After that, the wells were treated with a streptavidin-HRP conjugate, followed by TMB substrate to develop the color. Using a microplate reader (BioTek Synergy), the absorbance was measured at 450 nm after stopping the reaction with 2N sulfuric acid. We used a 4-parameter logistic (4-PL) model to fit standard curves and find cytokine concentrations.

#### 8-Hydroxy-2′-deoxyguanosine (8-OHdG)

We used a competitive ELISA kit (Abcam, UK) to measure oxidative DNA damage by quantifying 8-OHdG levels in serum. Standards and serum samples were added to wells precoated with an 8-OHdG conjugate, and the wells were incubated with an anti-8-OHdG primary antibody for 1 hour at 37 °C. After washing, a secondary antibody conjugated to HRP was added and incubated for 30 minutes. We used TMB as the substrate to generate the color, then stopped the reaction with 1 M HCl. We read the absorbance at 450 nm. Because it is a competitive assay, 8-OHdG concentrations were inversely related to optical density and were determined using a standard inhibition curve.

#### Total Antioxidant Capacity (TAC)

The ferric reducing antioxidant power (FRAP) assay was used to determine total antioxidant capacity. To make the FRAP working solution, we mixed acetate buffer (300 mM, pH 3.6), TPTZ (10 mM in 40 mM HCl), and FeCl□•6H□O (20 mM) in a 10:1:1 ratio. A 20 µL sample of serum was mixed with 180 µL of freshly made FRAP reagent and kept at 37 °C for 10 minutes. The ferrous form of the Fe^3^□-TPTZ complex was produced by reduction, resulting in a blue color at 593 nm. Using a FeSO□ standard curve, TAC values were found and given as µM Fe^2^□equivalents.

#### Hepatic and Renal Function Biomarkers

Standard colorimetric assay kits from Thermo Fisher Scientific (USA) were used to measure serum ALT, AST, urea, and creatinine levels. For ALT and AST, enzyme activity was measured using kinetic UV methods that monitored the rate of NADH oxidation. The absorbance was measured at 340 nm over 3 minutes. The urease–GLDH-coupled enzymatic method was used to determine urea concentration. In this method, the decrease in NADH absorbance at 340 nm was proportional to the amount of urea. The modified Jaffé reaction was used to measure creatinine. In this reaction, creatinine reacts with alkaline picrate to form an orange chromophore, which is measured at 520 nm. We ran each test three times and used kit-specific calibration standards to determine the results.

### 2.7. Histopathology

We collected liver and kidney tissues right after euthanasia and washed them gently in ice-cold phosphate-buffered saline (PBS) to get rid of any blood that was still there. To preserve cell structure, samples were fixed in 10% neutral buffered formalin (NBF; pH 7.2–7.4) for 24–48 hours at room temperature. After fixation, tissues were processed with an automated tissue processor (Leica Biosystems). This included sequential dehydration through graded ethanol series (70%, 80%, 90%, 95%, and 100%), clearing in xylene, and infiltration with melted paraffin wax. Using stainless-steel molds, the fixed tissues were then embedded in paraffin blocks. This ensured that the liver lobules and the renal cortex/medulla were in the correct positions. Using a rotary microtome (Leica RM2235), paraffin blocks were cut into four µm-thick sections. To remove wrinkles, the sections were floated in a 40 °C water bath and then placed on clean glass slides precoated with poly-L-lysine to aid tissue adherence. Before staining, the slides were dried at 37 °C overnight. To stain the sections, they were soaked in xylene for 2 × 5 minutes to remove the paraffin, then in distilled water to rehydrate. Standard hematoxylin and eosin (H&E) staining was done, and the slides were put in Harris hematoxylin for 5 minutes, rinsed under running tap water, differentiated in 1% acid alcohol, and blued in Scott’s tap water substitute. After that, the sections were counterstained with eosin Y for one to two minutes, dehydrated in a series of increasing concentrations of ethanol, cleared in xylene, and coverslipped with DPX mounting medium. A light microscope (Olympus BX53, Japan) was used to examine histology at 40× magnification.

### 2.8. Statistical Analysis

The data were presented as mean ± SEM, with each experimental group comprising eight animals (n = 8). One-way analysis of variance (ANOVA) was used to compare multiple groups. Then, Tukey’s multiple comparisons post hoc test was used to find the differences between groups. GraphPad Prism version 10 (GraphPad Software, USA) was used for all statistical analyses.

## 3. Results

### 3.1. Effect on Serum Cytokines

The LPS challenge raised serum TNF-α and IL-6 levels by about 129% and 131%, respectively, compared to control animals. Compared to LPS, Nutrishield® supplementation cut TNF-α levels by 51%, and compared to unencapsulated β-carotene, it cut them by 26%, bringing them back to levels close to baseline. In the same way, IL-6 levels were 53% lower than LPS and 24% lower than β-carotene. The levels of IL-10, which helps fight inflammation, increased by about 100% compared to LPS. This shows that the cytokine profile has changed significantly toward an anti-inflammatory profile. Giving LPS raised TNF-α and IL-6 levels a lot, but Nutrishield® brought cytokine levels back to normal (Table 1).

**Table 1.** Serum Cytokine Levels (pg/mL, mean ± SEM).

| GROUP | TNF-A | IL-6 | IL-10 |
| --- | --- | --- | --- |
| CONTROL | 62 ± 5 | 51 ± 4 | 39 ± 3 |
| LPS | 142 ± 8*** | 118 ± 7*** | 38 ± 2 |
| LPS + B-CAROTENE | 93 ± 6** | 74 ± 5** | 55 ± 3* |
| LPS + NUTRISHIELD® | 69 ± 4# | 56 ± 3# | 76 ± 3### |

\* $p < 0.05$ , \*\* $p < 0.01$ , \*\*\* $p < 0.001$ vs. Control; # $p < 0.01$ , ### $p < 0.001$ vs. LPS.

**Table 2.** Oxidative Stress and Antioxidant Indices.

| GROUP | 8-OHDG (NG/ML) | TAC (MM TROLOX EQ.) |
| --- | --- | --- |
| CONTROL | 2.1 ± 0.2 | 2.6 ± 0.1 |
| LPS | 5.4 ± 0.3*** | 1.1 ± 0.1*** |
| LPS + B-CAROTENE | 3.8 ± 0.2** | 1.52 ± 0.08** |
| LPS + NUTRISHIELD® | 2.3 ± 0.2# | 2.5 ± 0.1# |

\* $p < 0.05$ , \*\* $p < 0.01$ , \*\*\* $p < 0.001$ vs. Control; # $p < 0.01$ , ### $p < 0.001$ vs. LPS.

### 3.2. Oxidative Stress Biomarkers

LPS raised serum 8-OHdG levels by a lot (5.4 ± 0.3 ng/mL), but Nutrishield® treatment brought them down to 2.3 ± 0.2 ng/mL (p < 0.001). The Nutrishield® group had the highest TAC levels (2.5 ± 0.1 mM Trolox equivalents), which were 64% higher than the β-carotene group’s levels (1.52 ± 0.08 mM). After being exposed to LPS, serum 8-OHdG levels went up by about 157%, which shows that a lot of DNA damage was done by oxidative stress. Compared to LPS, Nutrishield® lowered 8-OHdG levels by 57%, and compared to unencapsulated β-carotene, it lowered levels by 39%. After giving LPS, the total antioxidant capacity went down by 58%, but Nutrishield® brought it back up to levels similar to those of the controls, which is a 127% increase compared to LPS and a 64% increase compared to β-carotene.

### 3.4. Serum Biochemistry and Organ Weights

Compared to animals that only got LPS, Nutrishield® brought hepatic enzyme levels back to normal (Table 3). LPS administration led to increases of 143% and 109% in ALT and AST levels, respectively, which means that the liver cells were damaged. Nutrishield® brought ALT and AST levels back to normal, lowering them by about 55% and 48% compared to LPS. Renal biomarkers exhibited a comparable trend, with creatinine and urea levels decreased by 42% and 36%, respectively, in relation to LPS-only subjects. Supplementary **Table S1-3** shows how nutrishield® β-Carotene and other supplements work as antioxidants in vitro.

**Table 3.** Serum Biochemistry and Organ Indices.

| PARAMETER | CONTROL | LPS | B-CAROTENE | NUTRISHIELD® |
| --- | --- | --- | --- | --- |
| ALT (U/L) | 42 ± 3 | 102 ± 5*** | 68 ± 4** | 46 ± 3# |
| AST (U/L) | 65 ± 5 | 136 ± 6*** | 92 ± 5** | 71 ± 4# |
| UREA (MG/DL) | 25 ± 2 | 42 ± 3** | 34 ± 3* | 27 ± 2# |
| CREATININE (MG/DL) | 0.42 ± 0.03 | 0.76 ± 0.05** | 0.59 ± 0.04* | 0.44 ± 0.03# |
| LIVER WT/BODY WT (%) | 4.2 ± 0.3 | 5.6 ± 0.4** | 4.8 ± 0.3* | 4.3 ± 0.2# |

\* $p < 0.05$ , \*\* $p < 0.01$ , \*\*\* $p < 0.001$ vs. Control; # $p < 0.01$ , ### $p < 0.001$ vs. LPS.

### 3.5. Histopathological Observations

H&E staining revealed that LPS induced significant vacuolation in hepatocytes, augmented the infiltration of inflammatory cells, and resulted in minimal cellular apoptosis. β-Carotene partially restored normal architecture, while livers treated with Nutrishield® exhibited nearly normal morphology with minimal inflammation (Figure 1A–C). LPS-treated mice exhibited kidney sections characterized by glomerular congestion and tubular necrosis. Mice that were treated with Nutrishield® did not have these problems (Figure 2A–C).

**Figure 1.**
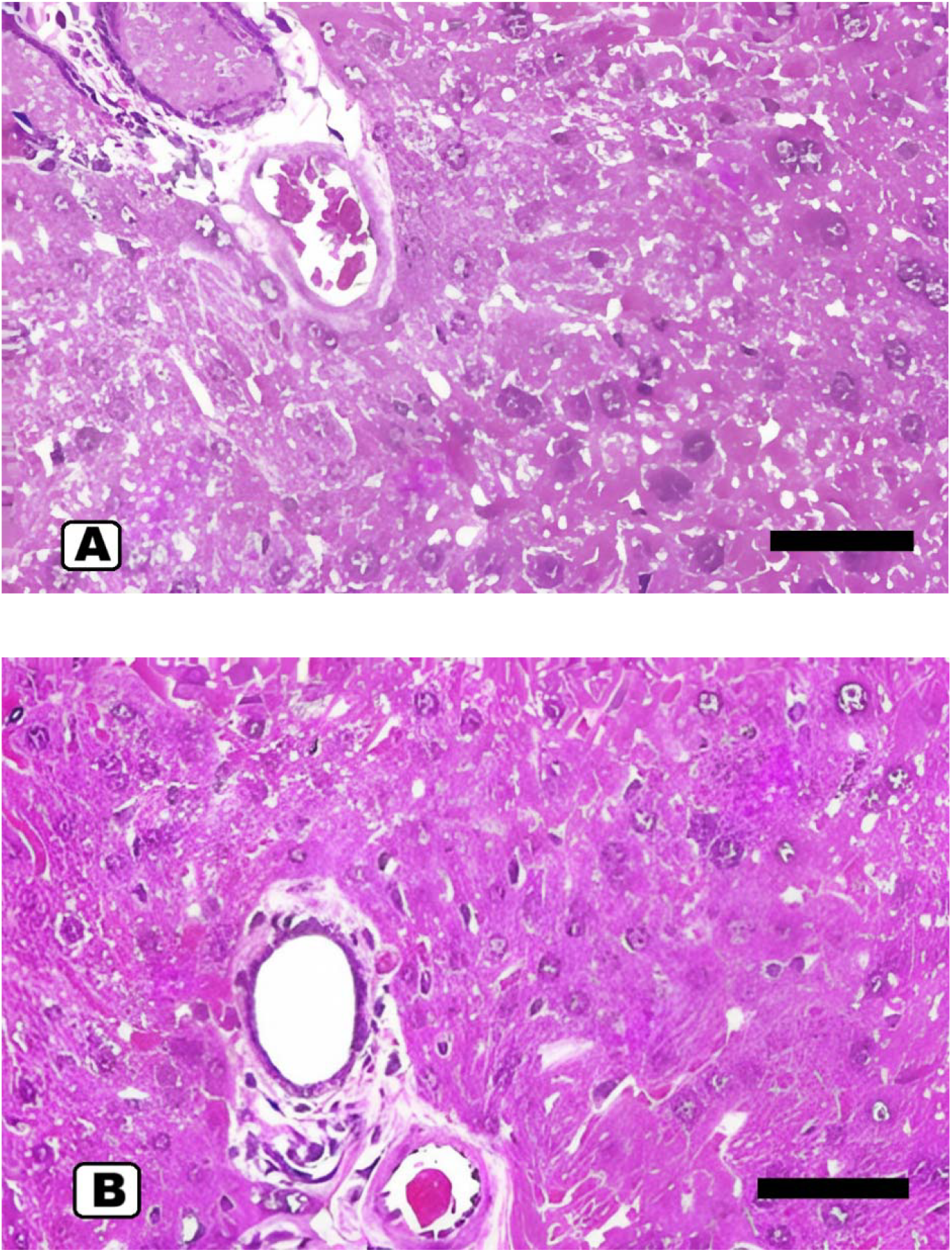
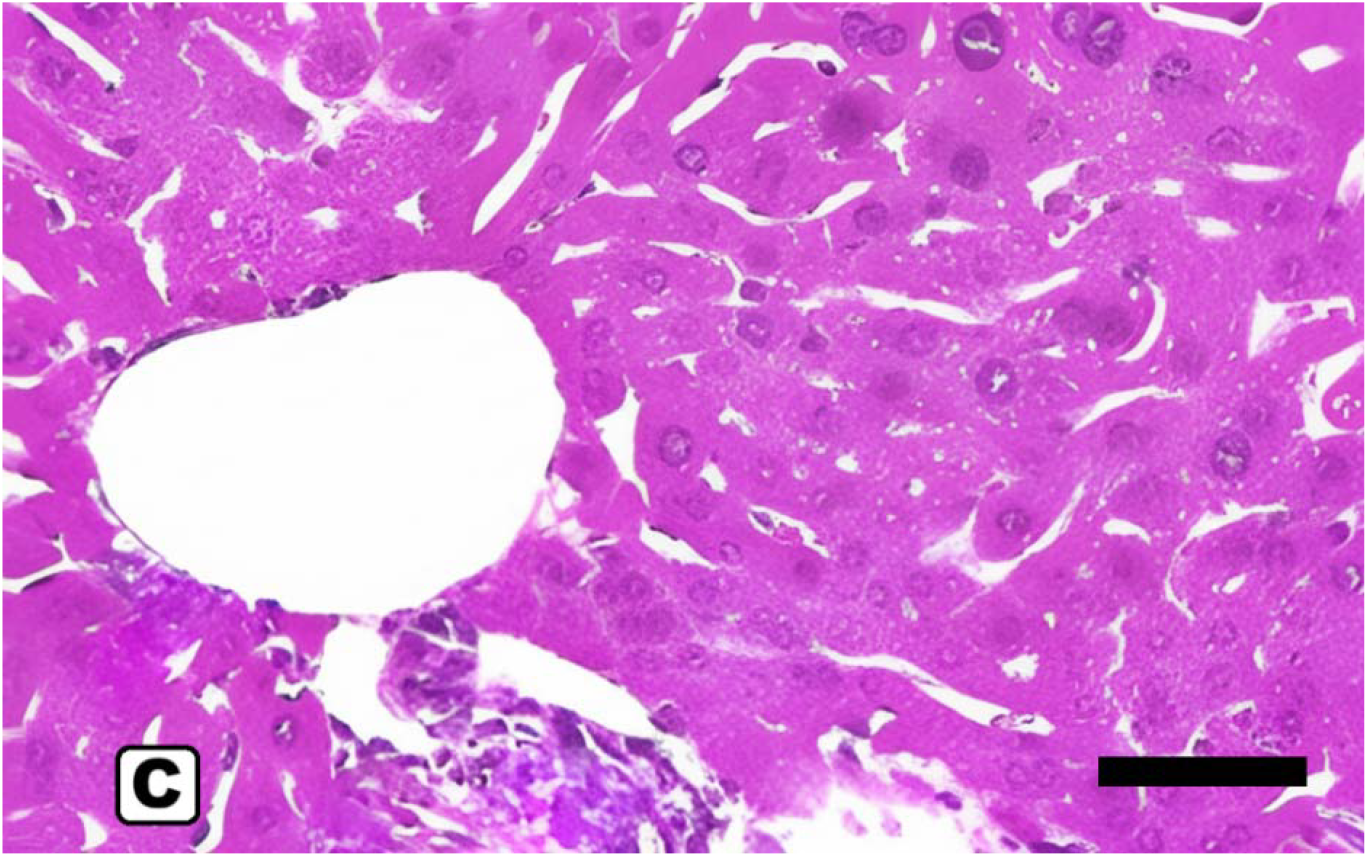
Liver histopathology (H&E, 40×): (A) LPS; (B) LPS + β-Carotene; (C) LPS + Nutrishield^®^. Scale bar, 50 μm.

**Figure 2.**
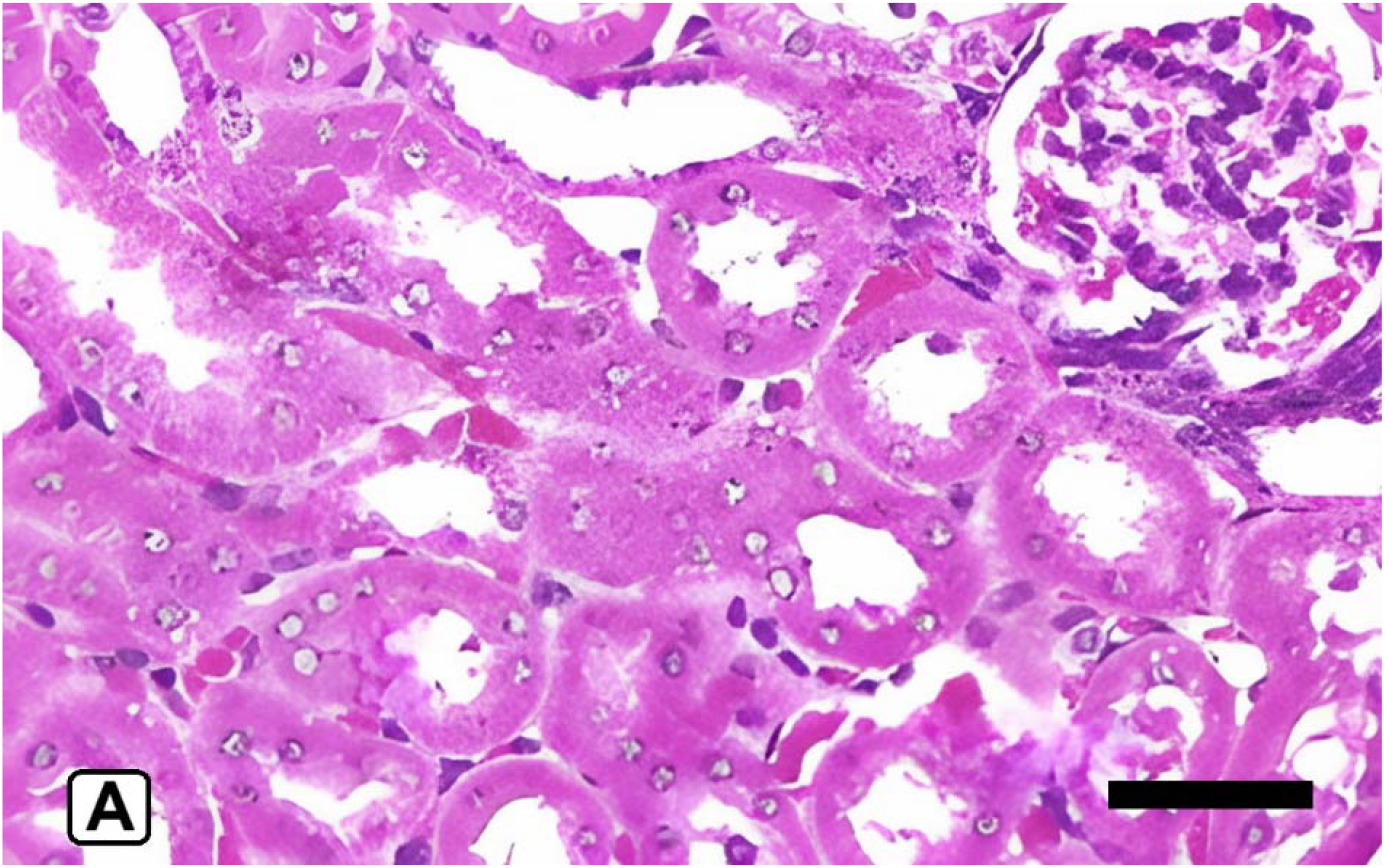
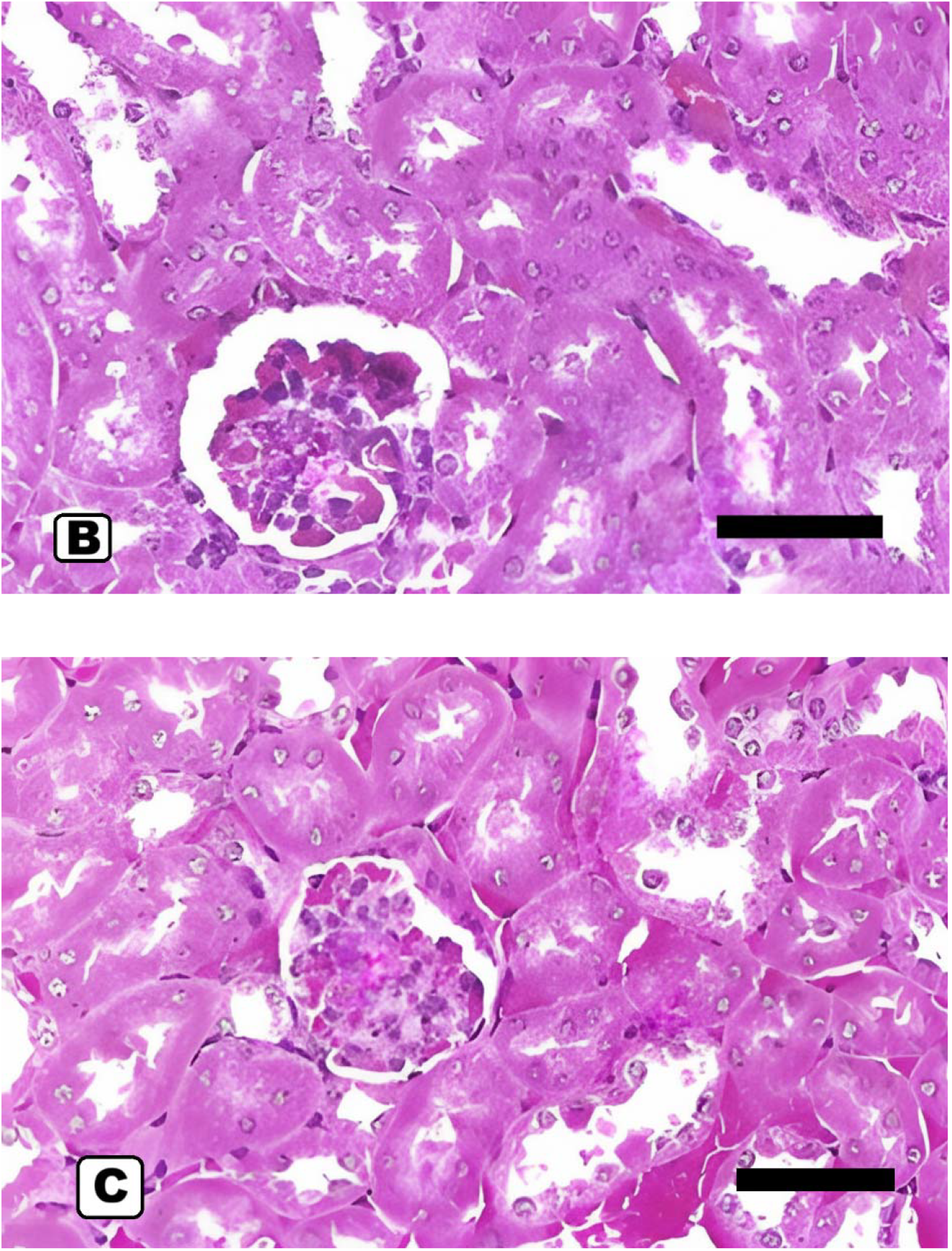
Kidney histopathology (H&E, 40×): (A) LPS; (B) LPS + β-Carotene; (C) LPS + Nutrishield^®^. Scale bar, 50 μm.

## 4. Discussion

This study demonstrated that microencapsulated Nutrishield® β-Carotene 20% TAB-S significantly mitigates LPS-induced systemic inflammation and oxidative stress compared to pure β-carotene. Animals administered the microencapsulated formulation demonstrated improved cytokine balance, enhanced antioxidant capacity, and significant hepatic and renal protection, thereby illustrating the functional advantages of the encapsulation technology. The bioactivity of β-carotene is closely associated with its capacity to neutralize reactive oxygen species, inhibit NF-κB activation, and regulate inflammatory signaling pathways (Li, Hong, & Zheng, 2019; Wu et al., 2023). When LPS triggers inflammation, TLR4 activation leads to the release of pro-inflammatory cytokines such as TNF-α, IL-6, and IL-1β, which worsen oxidative stress(Ciesielska, Matyjek, & Kwiatkowska, 2021). In this context, the administration of Nutrishield^®^ β-Carotene significantly normalized cytokine levels, indicating more effective suppression of LPS-induced inflammatory cascades than pure β-carotene.

These results align with prior studies indicating that β-carotene reduces IL-6 and TNF-α levels while increasing IL-10 production in LPS-challenged models (Li et al., 2019). The extent of cytokine modulation induced by Nutrishield^®^ β-Carotene exceeds that typically associated with unencapsulated β-carotene, underscoring the importance of delivery systems in maintaining carotenoid bioactivity. The starch-based microencapsulation matrix, aided by tocopherol stabilization, likely prevented β-carotene oxidation and made it easier for the body to absorb and distribute it. Prior research utilizing analogous encapsulation techniques—such as alginate–chitosan microcapsules, lipid nanocarriers, and cyclodextrin complexes—has demonstrated 2–5-fold increases in carotenoid absorption and biological efficacy (Bera, Mitra, & Singh, 2024; Soukoulis & Bohn, 2018).

The in vitro antioxidant assessment provides substantial mechanistic insight into the biological effects observed in the in vivo LPS-challenge model. Nutrishield^®^ β-Carotene consistently exhibited superior efficacy across all three antioxidant assays—DPPH, TPC, and ORAC—in comparison to all evaluated supplements. This multi-assay strength indicates that the antioxidant effects are not limited to specific assays; instead, they exhibit a general radical-neutralizing property unique to β-carotene. Due to its highly conjugated polyene structure, β-carotene has long been known to be a strong quencher of singlet oxygen and a stabilizer of lipid-free radicals. This mechanism elucidates the superior performance of Nutrishield^®^ β-Carotene compared to polyphenol-rich formulations: Phenolic antioxidants primarily act in water, while β-carotene acts in lipid compartments, where oxidative damage is most harmful, especially when inflammation is present, as in LPS-induced oxidative stress. The higher ORAC value supports this idea by showing that Nutrishield^®^ β-Carotene protects against oxidative damage caused by peroxyl radicals, one of the most biologically relevant radical species linked to tissue damage associated with inflammation.

Several other supplements tested in this study had moderate to high antioxidant values. The presence of polyphenols, anthocyanins, flavonoids, glutathione precursors, and herbal antioxidants is what makes them work. These molecules are known to donate hydrogen atoms or electrons to radicals, thereby making them more stable, as evidenced by their high DPPH and TPC readings. These formulations consistently exhibited lower antioxidant activity than Nutrishield^®^ β-Carotene, suggesting that although botanical compounds are effective in hydrophilic systems, they may lack the lipid-phase radical-quenching efficiency of β-carotene. Mineral-focused supplements like Calcium Complex and Calcium–Magnesium–Zinc–D3 had the lowest antioxidant values because they don’t have any antioxidants of their own. This serves as an internal negative control, confirming that the assays worked as expected and detected the presence or absence of active antioxidant ingredients. Moderate performance by multivitamin formulas indicates contributions from vitamins C and E and plant extracts; however, their efficacy did not reach the levels of either β-carotene or herbal products with higher polyphenol content.

Nutrishield^®^ β-Carotene strong antioxidant performance fits with what we already know about how carotenoids work together with other antioxidant systems. For instance, vitamin E and lycopene, both antioxidants often compared to β-carotene, have similar mechanisms for quenching singlet oxygen, stabilizing lipid peroxides, and maintaining redox balance in cells. Many studies show that carotenoids boost the immune system, reduce oxidative DNA damage, and alter the function of inflammatory signaling pathways (Kaulmann & Bohn, 2014; Milani, Basirnejad, Shahbazi, & Bolhassani, 2017). These similarities support the idea of bridging literature, leading to the conclusion that the antioxidant effects observed in vitro may also confer the protective effects observed in vivo. The in vitro antioxidant superiority of Nutrishield^®^ β-Carotene aligns with the biological effects demonstrated in the animal model, where β-carotene supplementation markedly diminished LPS-induced inflammation, reduced systemic oxidative stress, and maintained tissue architecture. Oxidative stress is a key factor in LPS-induced pathology because it increases cytokine release, lipid breakdown, and cellular damage. The in vitro results show that Nutrishield^®^ β-Carotene is a strong free radical neutralizer across multiple antioxidant mechanisms. This helps explain the anti-inflammatory, cytoprotective, and immunomodulatory effects seen in vivo. Overall, these results show that Nutrishield^®^ β-Carotene is a powerful antioxidant that can protect against oxidative stress in many ways. The comparator supplements are helpful in their own ways, such as supporting the urinary tract, improving vision, boosting metabolism, or maintaining multivitamin levels. However, they do not have the same range or level of antioxidant activity. This difference shows how valuable β-carotene is as an antioxidant that is embedded in membranes. It also supports the idea that it should be used in products intended to reduce inflammation and oxidative damage. The consistently high antioxidant performance observed across multiple Nutrishield^®^-containing supplements reinforces the platform functionality of the microencapsulation system. We anticipate that the Nutrishield® platform can be used for other vitamins, including vitamin B5. Nutrishield® can make vitamin B5 (pantothenic acid) work better by making it more stable and easier to absorb. It can also help keep the metabolic redox balance. Vitamin B5 is necessary for making energy in mitochondria, breaking down fatty acids, and controlling signals that cause inflammation. It is a precursor to coenzyme A (CoA), which is an important part. Controlled release from a starch-based encapsulating matrix could help the sodium-dependent multivitamin transporter (SMVT) not get too full too quickly. This would make it easier for the body to keep taking in the vitamins and keep plasma levels steady. Nutrishield^®^ platform demonstrated robust radical scavenging, reducing capacity, and peroxyl radical neutralization.

Better liver and kidney biomarkers support Nutrishield^®^ β-Carotene ability to protect against LPS-induced organ damage. These findings corroborate prior research indicating the hepatoprotective and nephroprotective effects of carotenoids in models of inflammatory, oxidative, and toxin-induced injuries (Elsayed et al., 2021). Studies using nanoencapsulated or emulsified carotenoids have demonstrated comparable organ-protective effects, attributable to enhanced retention, improved cellular uptake, and more effective ROS scavenging, underscoring the importance of formulation science in optimizing β-carotene efficacy (Dos Santos, Andrade, Flôres, & Rios, 2018; Gutiérrez et al., 2013; Zhang et al., 2016).

The data indicate that the microencapsulation of β-carotene in Nutrishield^®^ improves its therapeutic index by enhancing bioavailability, stabilizing the molecule against degradation, and facilitating sustained antioxidant activity. These traits make Nutrishield^®^ β-Carotene a promising nutraceutical for diseases like metabolic syndrome, nonalcoholic fatty liver disease (NAFLD), cardiovascular disease, inflammatory bowel disease, and age-related degenerative disorders that are caused by long-term inflammation and oxidative imbalance. Since β-carotene is known to support the immune system and protect the epithelial barrier, Nutrishield^®^ β-Carotene may also help with respiratory inflammation, immune modulation, and recovery after systemic inflammatory episodes. Future research and regulated human clinical trials would elucidate its translational potential and therapeutic significance.

## 5. Conclusion

In an LPS-induced murine inflammation model, Nutrishield^®^ β-Carotene 20% TAB-S, a starch-based microencapsulated β-carotene formulation, demonstrated significantly greater anti-inflammatory, antioxidant, and organ-protective effects than unencapsulated β-carotene. The enhanced effectiveness of Nutrishield^®^ β-Carotene is due to its improved physicochemical stability, controlled-release profile, and enhanced gastrointestinal tract absorption. All of these things make β-carotene more bioavailable in the body. Mice given Nutrishield^®^ β-Carotene showed a significant reduction in LPS-induced acute inflammation, as evidenced by reduced levels of pro-inflammatory cytokines (IL-6, TNF-α) in the blood and increased levels of the anti-inflammatory cytokine IL-10. The formulation also effectively protected against oxidative damage, as evidenced by lower levels of 8-OHdG in blood and higher total antioxidant capacity. This shows that it was very effective at protecting against injury caused by reactive oxygen species (ROS).

In conclusion, this preclinical study demonstrates that Nutrishield^®^ β-Carotene 20% TAB-S, a starch-based microencapsulated formulation, provides superior functional antioxidant and anti-inflammatory support compared with unencapsulated β-carotene in a murine model of systemic inflammation. The enhanced efficacy is attributable to improved stability, dispersion, and sustained biological activity conferred by the microencapsulation platform. These findings validate Nutrishield^®^ as a formulation-driven delivery system and support further translational evaluation.

## Acknowledgments

This work was funded by Singapore Ecommerce Centre Pte Ltd (Singapore), owner of the Nano Singapore brand. Grammarly (pro version) was used to refine the academic language. No AI-generated text was used directly in the final manuscript.

