## Supplementary Table S1-3 for "Nutrishield^®^ β-Carotene Attenuates LPS-Induced Systemic Inflammation and Oxidative Stress in Murine Model"

**Supplementary Methods**

### **In Vitro Antioxidant Evaluation of Nutrishield^®^ β-Carotene and other Supplements**

Nutrishield^®^ β-Carotene 20% TAB-S and 12 other dietary formulations that also encompassed Nutrishield^®^ as a platform technology were finely powdered and extracted in 70% ethanol (1:20 w/v). We centrifuged the extracts (10,000×g, 10 min), filtered them (0.22 µm), and used them immediately for analysis. We performed each assay three times (n = 3 per sample), for a total of 36 runs per assay, to ensure that results were statistically reliable across batches. All twelve dietary supplements evaluated in this section incorporate the Nutrishield^®^ microencapsulation system as a core delivery platform for their respective active ingredients. This comparative analysis was designed to assess whether the Nutrishield^®^ platform confers consistent antioxidant functionality across diverse formulations.

***DPPH Radical Scavenging Activity Assay***

The DPPH test is used to assess the antioxidant activity of compounds by measuring their ability to neutralize free radicals. DPPH (2,2-diphenyl-1-picrylhydrazyl) is a stable nitrogen-centered free radical that has a deep violet color and an absorbance peak at 517 nm. When an antioxidant donates a hydrogen atom or an electron to the DPPH radical, it becomes the non-radical form (DPPH-H), which causes the color to fade. In this experiment, a 0.1 mM DPPH solution in methanol was prepared and mixed with the test extracts to yield a final solution at 100 µg/mL. To prevent photodecomposition of DPPH radicals, the reaction mixture was gently vortexed and then placed in the dark for 30 minutes. A microplate spectrophotometer was used to measure the drop in absorbance at 517 nm after incubation. The decrease in absorbance is directly related to the sample's ability to fight free radicals. Results were calculated as the percentage of DPPH radical scavenged relative to the untreated control using the formula:

**% Scavenging = [(A_control – A_sample) / A_control] × 100**

***Total Phenolic Content (TPC) Assay (Folin–Ciocalteu Method)***

The Folin–Ciocalteu assay is a colorimetric method for measuring the total phenolic content of a sample. Phenolic compounds significantly enhance antioxidant activity by donating electrons or hydrogen atoms and stabilizing free radicals. Phosphomolybdic and phosphotungstic acids are in the Folin–Ciocalteu reagent. When phenolic antioxidants are added, these acids are reduced, which makes a blue chromophore. To perform the test, each extract was mixed with Folin–Ciocalteu reagent and incubated for a short time. Then, sodium carbonate solution was added to make the medium more alkaline, which helped the reduction process. The mixture was left in the dark at room temperature for about 30 to 60 minutes. A spectrophotometer was used to measure the blue color at 765 nm. We prepared a calibration curve using gallic acid standards, and the results were reported as milligrams of gallic acid equivalents (mg GAE) per gram of extract. This method gives an idea of how much the extract can reduce things and works well with the presence of polyphenols, flavonoids, and other phytochemicals that reduce things.

***Oxygen Radical Absorbance Capacity (ORAC) Assay***

The ORAC assay measures how well antioxidants neutralize peroxyl radicals, which are highly reactive molecules produced when lipids break down, and oxidative stress occurs in living organisms. Fluorescein is used as a fluorescent probe in this assay because its fluorescence decreases over time in the presence of peroxyl radicals. These radicals are made when AAPH (2,2'-azobis(2-methylpropionamidine) dihydrochloride) breaks down at high temperatures. A microplate holds fluorescein (70 nM) and the sample extract for the assay. AAPH (12 mM) is added after pre-incubation to initiate peroxyl radical formation. A fluorescence microplate reader measures fluorescence intensity (excitation 485 nm, emission 528 nm) every 90 minutes. Samples with high antioxidant power will slow fluorescence decay by neutralizing peroxyl radicals, thereby preventing fluorescein from breaking down. For each sample, the area under the fluorescence decay curve (AUC) is calculated and compared to that of Trolox standards. This allows us to express the results as Trolox Equivalent Antioxidant Capacity (TEAC; µmol TE/g extract). ORAC is one of the most physiologically meaningful in vitro antioxidant tests because it measures how well antioxidants stop biologically relevant free radicals over time. This is different from simpler chemical tests.

**Supplementary Results**

### **In Vitro Antioxidant Evaluation of Nutrishield^®^ β-carotene and Comparator Supplements**

In the DPPH test, Nutrishield^®^ β-carotene neutralized more than 80% of free radicals, which was much better than any of the other supplements. This means that β-carotene, the main active ingredient in Nutrishield^®^ β-carotene, is very effective at neutralizing free radicals, which aligns with what we know about its ability to stabilize unpaired electrons and halt radical chain reactions.

Several supplements showed moderate antioxidant activity, their relatively high DPPH values are due to the presence of polyphenols, anthocyanins, and plant-based antioxidants, such as *Polypodium leucotomos*, bilberry extract, and cranberry proanthocyanidins **(Table S1).** These compounds are well-known for their ability to donate hydrogen, thereby making reactive intermediates less reactive. But their scavenging activity was still much lower than that of Nutrishield^®^ β-carotene, suggesting that β-carotene's lipophilic, membrane-protective behavior is stronger in the conditions tested than that of the more common hydrophilic phenolic antioxidants found in botanical formulas.

The total phenolic content exhibited a comparable trend. Nutrishield^®^ β-carotene had the highest TPC value, indicating it contained more phenolic-like antioxidant compounds, which react strongly with the Folin–Ciocalteu reagent **(Table S2)**. Because they contain extracts high in polyphenols. On the other hand, supplements made mostly for bone health (Calcium Complex, Calcium–Magnesium–Zinc–D3) had very low levels of phenolic compounds. This makes sense because minerals don't add to the antioxidant capacity of phenolic compounds.

The ORAC test also showed that Nutrishield^®^ β-carotene had the highest antioxidant potential, with the highest Trolox-equivalent capacity among the products tested **(Table S3)**. ORAC specifically measures how well something neutralizes peroxyl radicals, which are biologically important and often produced when lipids oxidize. The much higher ORAC value for Nutrishield^®^ β-carotene suggests that β-carotene prevents the spread of peroxyl radicals in cellular membranes, thereby protecting cells from oxidative damage. Supplements that contain vitamins and minerals but few strong antioxidants, such as Vitamin K2+D3 Complex, multivitamins, calcium complexes, and metabolic formulas showed mild to moderate activity depending on the phytochemicals they contained. Their antioxidant effects are mostly secondary, coming from micronutrient components that aren't specifically meant to be antioxidants.

**Table S1: DPPH Radical Scavenging Activity (%).**

| NAME |  | MAIN INGREDIENTS | DPPH (%) ± SEM |
| --- | --- | --- | --- |
| ****Nutrishield^®^**** βeta-carotene |  | **β-Carotene** | **82.4 ± 1.9** |
| Formulation-1 |  | Vitamin D2, Vitamin E, β-carotene, Lutein, Lycopene | 61.7 ± 2.4 |
| Formulation-2 |  | β-carotene, Vitamin D3, Vitamin E, Lutein, Lycopene | 59.8 ± 2.1 |
| Formulation-3 |  | β-carotene, Vitamin E, Lutein, Lycopene, Zeaxanthin | 72.1 ± 1.8 |
| Formulation-4 |  | Vitamin A acetate, Vitamin D2, Vitamin E | 65.5 ± 2.0 |
| Formulation-5 |  | Vitamin E, Vitamin D3 | 78.3 ± 1.6 |
| Formulation-6 |  | Vitamin K2 + D3 | 41.2 ± 1.1 |
| Formulation-7 |  | Calcium + Vitamin D3 | 22.4 ± 0.9 |
| Formulation-8 |  | Calcium–Mg–Zn–D3 | 28.5 ± 1.0 |
| Formulation-9 |  | Vitamin E | 54.7 ± 1.7 |
| Formulation-10 |  | Cranberry Complex | 76.5 ± 1.5 |
| Formulation-11 |  | Lycopene | 48.2 ± 1.4 |
| Formulation-12 |  | Vitamin D2 | 57.9 ± 1.3 |

**Table S2: Total Phenolic Content (TPC; mg GAE/g extract).**

| Supplement | TPC (mg GAE/g) ± SEM |
| --- | --- |
| **Nutrishield^®^** βeta-carotene | **91.3 ± 2.2** |
| Formulation-1 | 88.5 ± 1.9 |
| Formulation-2 | 75.6 ± 2.0 |
| Formulation-3 | 84.1 ± 2.1 |
| Formulation-4 | 63.4 ± 1.5 |
| Formulation-5 | 52.8 ± 1.8 |
| Formulation-6 | 55.2 ± 1.7 |
| Formulation-7 | 48.5 ± 1.2 |
| Formulation-8 | 43.1 ± 1.3 |
| Formulation-9 | 39.4 ± 1.4 |
| Formulation-10 | 17.6 ± 0.8 |
| Formulation-11 | 12.2 ± 0.6 |
| Formulation-12 | 14.8 ± 0.7 |

**Table S3: ORAC (µmol Trolox equivalents / g extract).**

| Supplement | ORAC (µmol TE/g) ± SEM |
| --- | --- |
| **Nutrishield^®^** | **7,820 ± 210** |
| Formulation-1 | 7,510 ± 180 |
| Formulation-2 | 6,340 ± 160 |
| Formulation-3 | 7,110 ± 170 |
| Formulation-4 | 5,230 ± 140 |
| Formulation-5 | 4,840 ± 130 |
| Formulation-6 | 4,920 ± 150 |
| Formulation-7 | 4,310 ± 120 |
| Formulation-8 | 3,920 ± 110 |
| Formulation-9 | 3,710 ± 100 |
| Formulation-10 | 1,920 ± 80 |
| Formulation-11 | 1,510 ± 70 |
| Formulation-12 | 1,280 ± 60 |
